# Kalium rhodopsins: Natural light-gated potassium channels

**DOI:** 10.1101/2021.09.17.460684

**Authors:** Elena G. Govorunova, Yueyang Gou, Oleg A. Sineshchekov, Hai Li, Yumei Wang, Leonid S. Brown, Mingshan Xue, John L. Spudich

**Affiliations:** Center for Membrane Biology, Department of Biochemistry & Molecular Biology, The University of Texas Health Science Center at Houston McGovern Medical School; Houston, TX 77030, USA; Department of Neuroscience, Baylor College of Medicine; Houston, TX 77030, USA; The Cain Foundation Laboratories, Jan and Dan Duncan Neurological Research Institute at Texas Children’s Hospital; Houston, Texas 77030, USA; Department of Physics and Biophysics Interdepartmental Group, University of Guelph; Guelph, Ontario N1G 2W1, Canada; Department of Molecular & Human Genetics, Baylor College of Medicine; Houston, TX 77030, USA

## Abstract

We report a family of K^+^ channels, kalium channelrhodopsins (KCRs) from a fungus-like protist. Previously known potassium channels, widespread and mainly ligand- or voltage-gated, share a conserved pore-forming domain and K^+^-selectivity filter. KCRs differ in that they are light-gated and they have independently evolved an alternative K^+^ selectivity mechanism. The KCRs are potent, highly selective of K^+^ over Na^+^, and open in less than 1 millisecond following photoactivation. Their permeability ratio P_K_/P_Na_ of ∼ 20 make KCRs powerful hyperpolarizing tools that suppress excitable cell firing upon illumination, demonstrated here in mouse cortical neurons. KCRs enable specific optogenetic photocontrol of K^+^ gradients promising for the study and potential treatment of potassium channelopathies such as epilepsy, Parkinson’s disease, and long-QT syndrome and other cardiac arrhythmias.

**One-Sentence Summary:** Potassium-selective channelrhodopsins long-sought for optogenetic research and therapy of neurological and cardiac diseases.

## Main Text

Potassium (K^+^) channels, ubiquitously found in all domains of life are easily recognized by their highly conserved “K^+^ channel signature sequence” (*1-4*) that encodes a K^+^-selectivity filter that strongly favors conductance of K^+^ over Na^+^. We report here a type of K^+^ channel that defines a unique family in that its members (i) completely lack the signature sequence of previously known K^+^ channels, and (ii) unlike prior K^+^ channels, they are channelrhodopsins, retinylidene proteins gated by light.

Channelrhodopsins (ChRs) are light-gated ion channels first discovered in the model chlorophyte alga *Chlamydomonas reinhardtii* that serve as membrane-depolarizing photoreceptors in phototactic protists (*5-7*). They are used to manipulate the membrane potential of excitable animal cells with light (optogenetics) (*8*). Cation conductive ChRs (CCRs) are primarily proton channels, some of which also conduct mono- and divalent metal cations (*7, 9-11*). Photoactivation of CCRs depolarizes the neuronal membrane by Na^+^ and H^+^ influx, and stimulates spiking (*12*). Anion conductive ChRs (ACRs) conduct halides and nitrate (*13*). Their photoactivation hyperpolarizes or depolarizes the neuronal membrane depending on the electrochemical gradient of Cl^-^, and inhibits or stimulates spiking, respectively (*14, 15*).

The electrochemical gradient of K^+^ favors membrane hyperpolarization in neurons, which has stimulated efforts to engineer light-gated K^+^ channels to be used as neuronal silencing tools. The K^+^/Na^+^ permeability ratio (P_K_/P_Na_) of *C. reinhardtii* ChR2 (*Cr*ChR2), the most used optogenetic variant, is 0.3-0.5 (*7, 16, 17*). Some mutations increased the P_K_/P_Na_ ratio of *Cr*ChR2, but no more than twice (*17*). Alternatively, K^+^ channels have been modified to become photosensitive by the addition of synthetic photoswitches or photoactive protein domains. Recent results were obtained from fusing the photoreceptor LOV2 domain with a K^+^ channel (*18*), and indirect control by co-expressing a photosensitive adenylyl cyclase and a cAMP-gated K^+^ channel (*19, 20*). Both approaches are promising for some applications, but are limited by the slow kinetics of the LOV2/channel chimera (BLINK2) and possible cAMP-induced side effects. Here we show that two ChRs from *Hyphochytrium catenoides*, which we named *Hc*KCR1 and *Hc*KCR2 (for *<u>H</u>yphochytrium <u>c</u>atenoides* <u>k</u>alium <u>c</u>hannelrhodopsins), are highly specific, robust light-gated K^+^ channels that open on the submillisecond time scale.

*H. catenoides* is a fungus-like heterotrophic organism from the stramenopile class Hyphochytriomycetes, the genome of which has been sequenced (*21*). Two predicted *H. catenoides* proteins show homology to bacteriorhodopsin-like CCRs (BCCRs) from cryptophyte algae (fig. S1). In contrast to other known ChRs, BCCRs contain Asp residues in the positions of the photoactive site (retinylidene Schiff base) proton donor and acceptor of bacteriorhodopsin (Asp85 and Asp96, respectively) (*11*), conserved in the *H. catenoides* homologs (fig. S2). A recent high-resolution structure of a *Rhodomonas lens* BCCR known as ChRmine suggested trimeric organization with a conductive pore between subunits (*22*). However, out of the residues implicated in the trimer formation in ChRmine, only Glu68 is conserved in the *H. catenoides* homologs (fig. S2).

We synthesized mammalian-codon adapted versions of the polynucleotides encoding the heptahelical transmembrane (rhodopsin) domains and expressed them in human embryonic kidney (HEK293) cells as mCherry fusions. Both homologs were photoactive; the action spectra of their photocurrents are shown in Fig. 1A. Following a historical convention, we assigned the number 1 to the more red-shifted paralog (spectral maximum 540 nm), and the number 2, to the other (spectral maximum 490 nm).

**Fig. 1.**
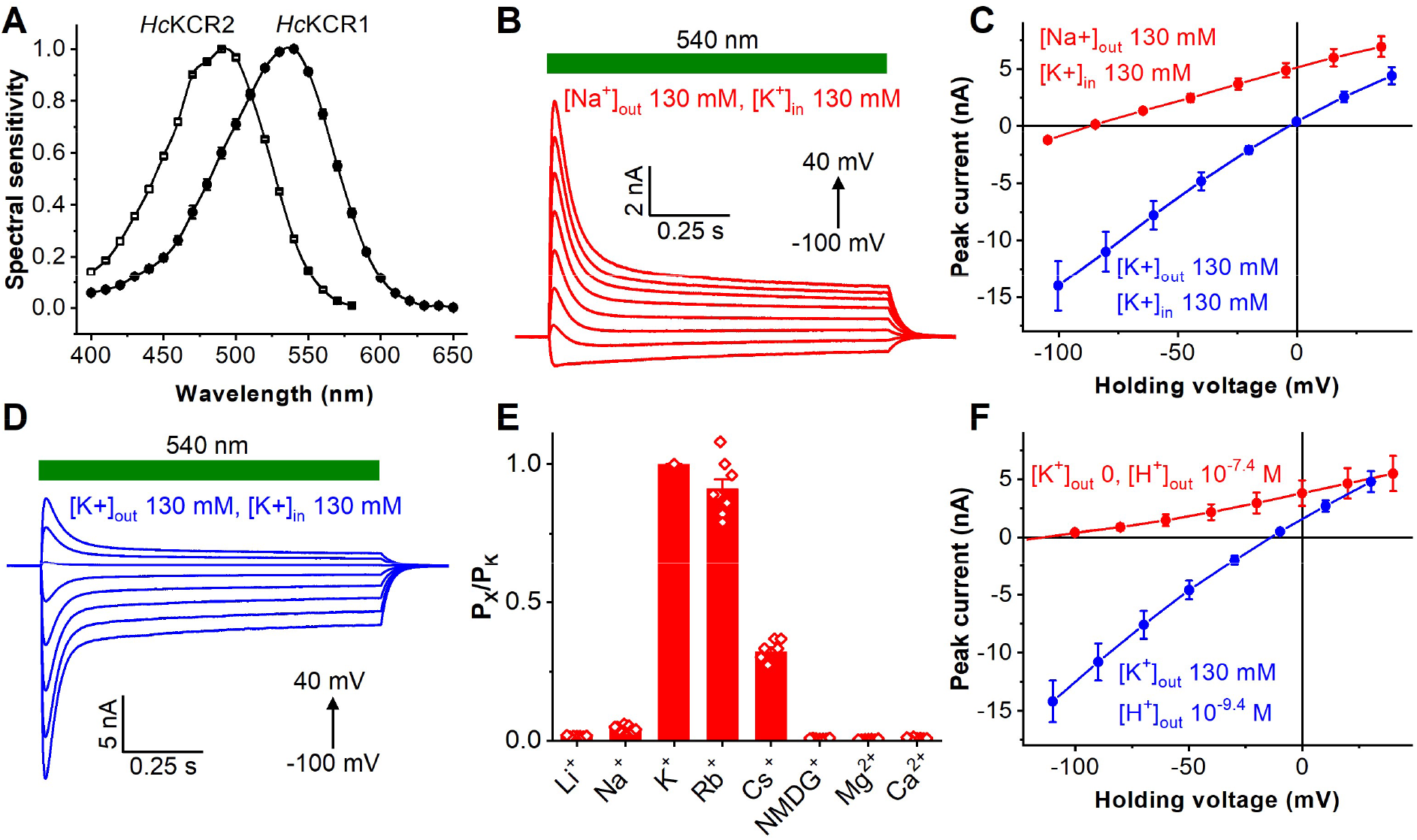
KCR photocurrents evoked by pulses of continuous light. (**A**) The action spectra of photocurrents (mean ± sem, n = 7 scans). (**B** and **D**) Photocurrents traces recorded from *Hc*KCR1 in response to 1-s light pulses upon 20-mV voltage increments. (**C** and **F**) The *IE* curves (mean ± sem, n = 7 cells). (**E**) The permeability ratios (P_X_/P_K_); mean ± sd; the diamonds, the data from individual cells.

A series of photocurrents generated by *Hc*KCR1 in response to pulses of continuous light under physiological ionic conditions (130 mM K^+^ in the pipette and 130 mM Na^+^ in the bath, both pH 7.4, for full solution compositions see table S1), is shown in Fig. 1B. The nearly linear current-voltage relationships (*IE* curves) showed a steep slope with a reversal potential (*E*_*rev*_) of - 85 ± 2 mV (mean ± sem, n = 10 cells) (Fig. 1C, red). Such behavior has not been observed in any previously tested ChRs, and could only be explained by selectivity for K^+^ over Na^+^, as the concentration of Cl^-^ was nearly identical on the two sides of the membrane. This conclusion was confirmed by the shift of the *E*_*rev*_ to -3 ± 1 mV (mean ± sem, n = 7 cells) measured upon complete replacement of Na^+^ in the bath with K^+^ (Figs. 1C and D, blue). The corresponding results for *Hc*KCR2 are shown in fig. S3. The P_K_/P_Na_ permeability ratio of *Hc*KCR1 calculated using the modified Goldman-Hodgkin-Katz (GHK) voltage equation (*23*) was 23, 60 times greater than that of *Cr*ChR2. The P_K_/P_Na_ value of *Hc*KCR2 was 17.

As *Hc*KCR1 exhibited a more red-shifted spectrum, larger current amplitude and higher selectivity for K^+^ than *Hc*KCR2, we chose this channel for a more detailed characterization. Using a similar procedure as for Na^+^, we determined the P_X_/P_K_ ratios for other metal cations and *N*-methyl-*D*-glucamine (NMDG^+^). Representative series of photocurrent traces and mean *IE* curves are shown in fig. S4, and the P_X_/P_K_ values, in Fig. 1E. The permeability sequence of *Hc*KCR1 was K^+^ > Rb^+^ > Cs^+^ > Na^+^ > Li^+^ > NMDG^+^ ≅ Mg^2+^ ≅ Ca^2+^ (Eisenman sequence IV), the same as that of most voltage- and ligand-gated K^+^ channels (*24*). To estimate an upper limit of the P_H_/P_K_ ratio, we measured the *E*_*rev*_ shift between the bath containing K^+^ at pH 9.4 and non-permeable NMDG^+^ at pH 7.4 (Fig. 2F, red). The P_H_/P_K_ calculated from this experiment was ∼3 × 10^4^, ∼80 times lower than that of *Cr*ChR2 (*7, 16, 17*).

**Fig. 2.**
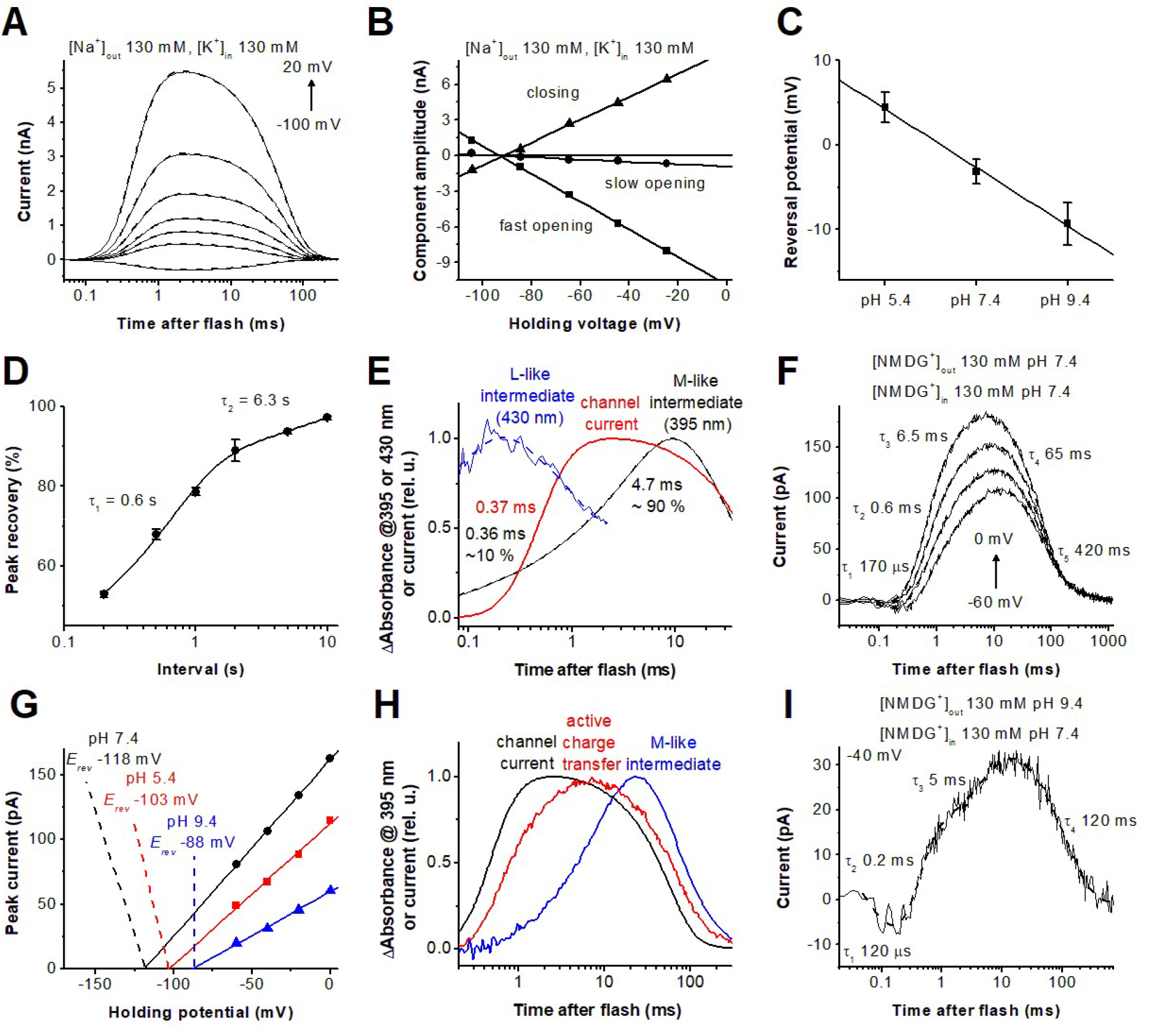
Photocurrents and photochemical conversion upon single quantum excitation. (**A**) Photocurrent traces (thin solid lines) recorded from *Hc*KCR1 in response to laser flashes upon 20-mV voltage increments under the indicated ionic conditions superimposed with their multiexponential approximations (dashed lines). (**B**) Voltage dependence of the three kinetic components of channel currents. (**C**) Dependence of the channel current *E*_*rev*_ on bath pH (mean ± sem, n = 3-8 cells). (**D**) Time course of the peak current recovery (mean ± sem, n = 5 cells). (**E**) Transient absorbance changes (blue and black) and channel current (red). (**F**) Photocurrent traces in the absence of permeant metal cations at bath pH 7.4. (**G**) Voltage dependence of active current at different bath pH. (**H**) Transient absorbance changes at 400 nm (blue), compared to active and channel currents (red and black, respectively). (**I**) Photocurrent traces in the absence of permeant metal cations at bath pH 9.4.

We used 6-ns laser flash excitation to follow the kinetics of channel gating, probe for active charge movements and eliminate effects of second photon absorption. Regardless of ionic gradients, channel currents could be fit with three exponentials (Fig. 2A and fig. S5A). Channel opening was biphasic, as in *Gt*ACR1 (*25*) and *Cr*ChR2 (*26*), but channel closing was monophasic, unlike these two ChRs. The fast opening accelerated, and the slow opening slowed upon depolarization (fig. S5B). The *E*_*rev*_ of the amplitudes of the three kinetics components were the same in all experimental conditions (Fig. 2B and fig. S5C). Our interpretation is that the relative permeability for different cations does not change during opening and closing of the channel. With equal concentrations of K^+^ on both sides of the membrane the *E*_*rev*_ depended on bath pH (Fig. 2C). However, the difference between pH 5.4 and 9.4 was only ∼15 mV, which confirmed the low P_H_/P_K_ of *Hc*KCR1 relative to earlier known CCRs.

The photocurrent decayed to the noise level in < 200 ms after the flash. To estimate the time for complete dark recovery we applied a series of laser flashes with progressively shorter intervals between them (fig. S6). The recovery was biphasic with τ = 0.6 and 6.6 s (Fig. 2D). We observed multiphasic recovery with similar τ values by flash photolysis in both detergent-purified pigment and *Pichia* membranes (fig. S7).

Absorbance at 430 nm dropped concomitant with an increase at 395 nm, which indicated that these wavelengths were characteristic of an L-like and an M-like intermediate, respectively (Fig. 2E). Opening of the channel occurred upon transition from the late L to the early M intermediate. The M rise was biphasic, and τ of the fast component was close to that of channel opening unlike both *Gt*ACR1, in which channel opening takes place in the L state (*25, 27*), and “classical” chlorophyte CCRs, in which M rise (i.e. deprotonation of the retinylidene Schiff base) precedes channel opening (*28, 29*).

In previously characterized BCCRs, channel conductance was found to be tightly coupled to active vectorial transport of protons (*11*). Upon substitution of non-permeable NMDG^+^ for K^+^ and Na^+^, photocurrents several times slower than channel current were recorded (Fig. 2F). Their voltage dependence crossed the X axis at very negative values, characteristic of active charge movement (Fig. 2G). These values remained close even when the difference in the H^+^ gradient was varied over four orders of magnitude (i.e. 240 mV), suggesting that the photoactive site was barely accessible to protons from outside, as expected from the relatively low proton permeability of *Hc*KCR1.

The initial unresolved negative component of charge movement is a typical reflection of retinal isomerization, reported earlier in other ChRs (*29*). Rise of positive photocurrent was biphasic with τ values similar to those of components of biphasic M rise, indicating active proton transfer from the Schiff base to an outwardly located acceptor (Fig. 2E, F). However, the peak of the current was reached before that of M accumulation, and the current decayed in the time window of M decrease (Fig. 2H). This observation suggests that reprotonation of the Schiff base at least partially takes place inside the photoactive site from the initially protonated acceptor, and there is no actual proton pumping across the membrane. In some recordings the decay of active charge movement could be resolved in two exponentials with τ = 65 and 420 ms (Fig. 2F). However, in most cases the two decay components were merged into a single one with τ ∼120 ms (Fig. 2I and fig. S8). At an increased outwardly directed H^+^ gradient the rates of both positive currents only slightly increased (Fig. 2I), which confirmed the isolation of the photoactive site from the outside medium. Such isolation is unusual for H^+^-pumping rhodopsins and may be related to the high K^+^ selectivity of the *Hc*KCR1 channel.

We tested whether *Hc*KCR1 can be used to suppress excitable cell firing. *Hc*KCR1 fused with EYFP and tdTomato were expressed in layer 2/3 pyramidal neurons of the mouse somatosensory cortex by *in utero* electroporation (Fig. 3A). We prepared acute brain slices from 2–4-week-old mice and performed whole-cell voltage clamp recording from *Hc*KCR1-expressing neurons with 142 mM K^+^ in the pipette and 2.5 mM K^+^ in the bath (for full solution compositions see Methods). In response to 1-s pulses of green light, *Hc*KCR1 generated robust photocurrents (Fig. 3B) that recovered quickly in the dark (fig. S9). The decay of photocurrents was best fit by two exponentials with τ_1_ = 40 ms (Fig. 3C) and τ_2_ = 0.6 ± 0.1 s at -45 mV and 0.5 ± 0.2 at -85 mV (mean ± sem, n = 8 cells). The ratio of steady-state to peak photocurrents increased when the membrane was hyperpolarized (Fig. 3D). The *IE* curves showed a reversal potential of -63 mV for peak photocurrents and -56 mV for steady-state photocurrents (Fig. 3E-G), indicating that channel states formed upon absorption of a second photon under continuous light stimulation alter the relative permeability for cations. We next performed current clamp recordings to test *Hc*KCR1 as a neuronal silencing tool. Photoactivation of *Hc*KCR1 instantly and persistently inhibited all action potentials induced by current injections (Fig. 3H, I), demonstating that *Hc*KCR1 is a potent optogenetic silencer.

**Fig. 3.**
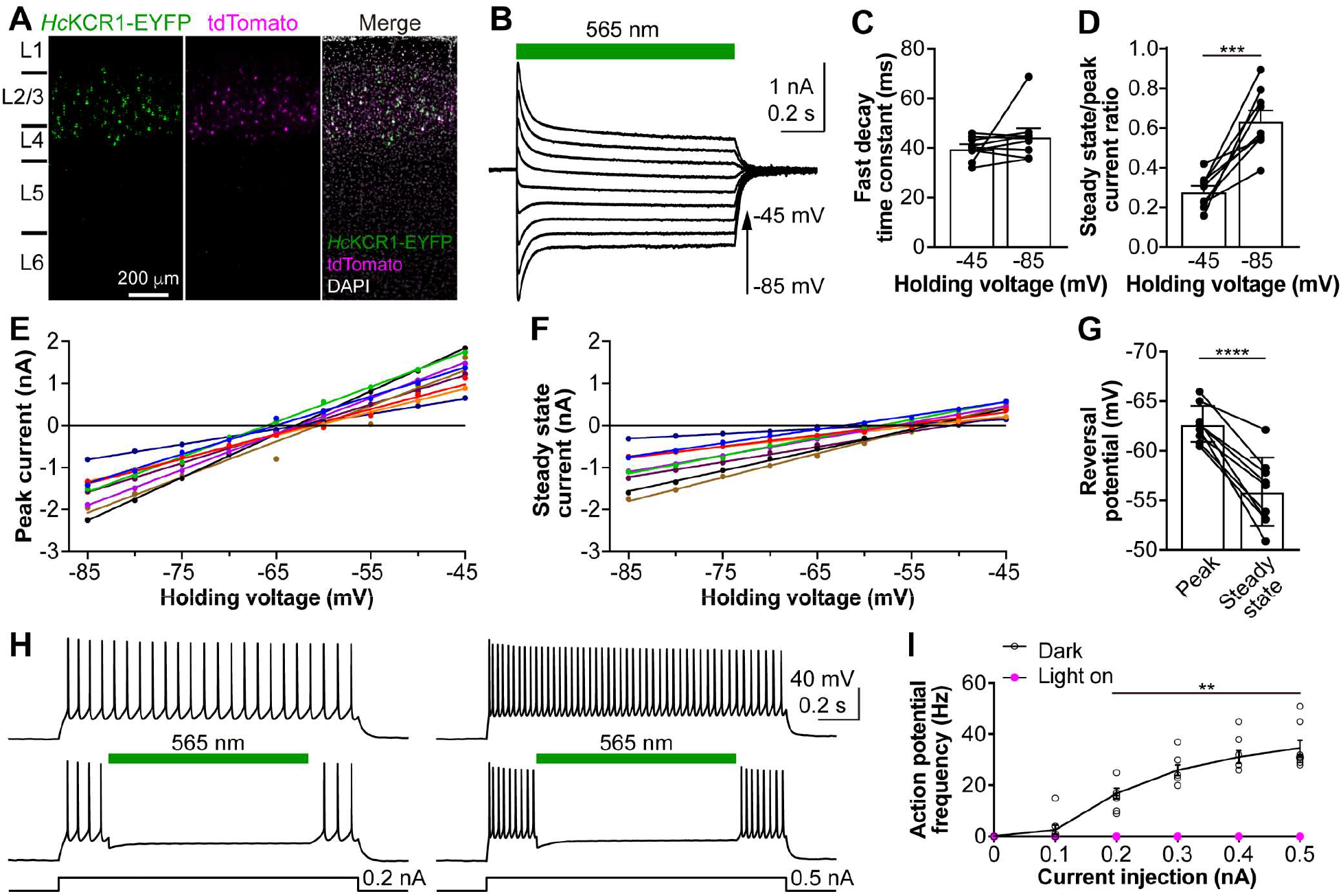
Photoactivation of *Hc*KCR1 in neurons generates robust photocurrents and efficiently suppresses neuronal firing. (**A**) Fluorescent images of a cortical slice showing *Hc*KCR1-EYFP and tdTomato expression in layer 2/3 neurons. Cortical layers were identified by DAPI staining. L, layer. (**B**) Photocurrents traces of a *Hc*KCR1-expressing neuron in response to a 1-s 565 nm light pulse at holding voltages increased in 5-mV steps. (**C**) The fast time constant of photocurrent decay at indicated voltages. (**D**) Ratios of steady-state to peak photocurrents. (**E, F**) *IE* curves of peak (E) and steady state (F) photocurrent of individual neurons indicated by different colors. (**G**) Reversal potentials calculated from the data in E and F. (**H**) Membrane voltage traces of a *Hc*KCR1-expressing neuron in response to 0.2 (left) and 0.5 nA (right) current injections without (top) and with (bottom) 565 nm light pulses. (**I**) The frequencies of action potentials evoked by different current injections with (magenta) and without (black) photoactivation. Data in C, D, G and I are expressed as mean ± sem, n = 8 cells; ** *P* < 0.01, *** *P* < 0.001, **** *P* < 0.0001.

The discovery of KCRs provides an alternative mechanism for K^+^ selection and our studies lay the basis for its elucidation. KCRs also expand the optogenetic toolbox with a natural K^+^-selective tool that benefits from the high efficiency provided by evolution, enabling direct, rapid, and potent photocontrol of K^+^ transmembrane gradients.

## Author contributions

Conceptualization: EGG, OAS, LSB, MX, JLS

Methodology: EGG, OAS, LSB, MX, JLS

Investigation: EGG, YG, OAS, HL, YW, LSB

Visualization: EGG, YG, OAS

Funding acquisition: LSB, MX, JLS

Project administration: JLS Supervision: JLS

Writing – original draft: EGG, YG, OAS, MX, JLS

Writing – review & editing: EGG, YG, OAS, HL, YW, LSB, MX, JLS

## Data and materials availability

All data and materials are available to any researcher upon a reasonable request, pending materials transfer agreements (MTAs). The sequences of *Hc*KCR1 and *Hc*KCR2 expression constructs are available from GenBank (accession numbers MZ826862 and MZ826861, respectively).

## Supplementary Material

Materials and Methods

Figs. S1 to S9

Table S1

References (*30-36*)

## References and Notes

1. R. MacKinnon, Potassium channels. FEBS Lett. 555, 62–65 (2003).

2. P. Enyedi, G. Czirjak, Molecular background of leak K^+^ currents: two-pore domain potassium channels. Physiol. Rev. 90, 559–605 (2010).

3. T. D. Mackie, J. L. Brodsky, Investigating potassium channels in budding yeast: A genetic sandbox. Genetics 209, 637–650 (2018).

4. A. Mironenko, U. Zachariae, B. L. de Groot, W. Kopec, The persistent question of potassium channel permeation mechanisms. J. Mol. Biol. 433, 167002 (2021).

5. O. A. Sineshchekov, K.-H. Jung, J. L. Spudich, Two rhodopsins mediate phototaxis to low- and high-intensity light in Chlamydomonas reinhardtii. Proc. Natl. Acad. Sci. USA 99, 8689–8694 (2002).

6. G. Nagel et al., Channelrhodopsin-1: a light-gated proton channel in green algae. Science 296, 2395–2398 (2002).

7. G. Nagel et al., Channelrhodopsin-2, a directly light-gated cation-selective membrane channel. Proc. Natl. Acad. Sci. USA 100, 13940–13945 (2003).

8. K. Deisseroth, Optogenetics. Nat. Methods 8, 26–29 (2011).

9. E. G. Govorunova, E. N. Spudich, C. E. Lane, O. A. Sineshchekov, J. L. Spudich, New channelrhodopsin with a red-shifted spectrum and rapid kinetics from Mesostigma viride. mBio 2, e00115–00111 (2011).

10. E. G. Govorunova, O. A. Sineshchekov, H. Li, R. Janz, J. L. Spudich, Characterization of a highly efficient blue-shifted channelrhodopsin from the marine alga Platymonas subcordiformis. J. Biol. Chem. 288, 29911–29922 (2013).

11. O. A. Sineshchekov, E. G. Govorunova, H. Li, J. L. Spudich, Bacteriorhodopsin-like channelrhodopsins: Alternative mechanism for control of cation conductance. Proc. Natl. Acad. Sci. USA 114, E9512–E9519 (2017).

12. E. S. Boyden, F. Zhang, E. Bamberg, G. Nagel, K. Deisseroth, Millisecond-timescale, genetically targeted optical control of neural activity. Nat. Neurosci. 8, 1263–1268 (2005).

13. E. G. Govorunova, O. A. Sineshchekov, X. Liu, R. Janz, J. L. Spudich, Natural light-gated anion channels: A family of microbial rhodopsins for advanced optogenetics. Science 349, 647–650 (2015).

14. M. Mahn, M. Prigge, S. Ron, R. Levy, O. Yizhar, Biophysical constraints of optogenetic inhibition at presynaptic terminals. Nat. Neurosci. 19, 554–556 (2016).

15. J. E. Messier, H. Chen, Z. L. Cai, M. Xue, Targeting light-gated chloride channels to neuronal somatodendritic domain reduces their excitatory effect in the axon. eLife 7, e38506 (2018).

16. J. Y. Lin, M. Z. Lin, P. Steinbach, R. Y. Tsien, Characterization of engineered channelrhodopsin variants with improved properties and kinetics. Biophys. J. 96, 1803–1814 (2009).

17. R. Richards, R. E. Dempski, Re-introduction of transmembrane serine residues reduce the minimum pore diameter of channelrhodopsin-2. PLoS One 7, e50018 (2012).

18. L. Alberio et al., A light-gated potassium channel for sustained neuronal inhibition. Nat Methods 15, 969–976 (2018).

19. S. Beck et al., Synthetic light-activated ion channels for optogenetic activation and inhibition. Front. Neurosci. 12, 643 (2018).

20. Y. A. Bernal Sierra et al., Potassium channel-based optogenetic silencing. Nat. Commun. 9, 4611 (2018).

21. G. Leonard et al., Comparative genomic analysis of the ‘pseudofungus’ Hyphochytrium catenoides. Open. Biol. 8, (2018).

22. K. E. Kishi et al., Structural basis for channel conduction in the pump-like channelrhodopsin ChRmine. bioRxiv, https://www.biorxiv.org/content/10.1101/2021.1108.1115.456392v456391 (2021).

23. B. Hille, Ion channels of excitable membranes. (Sinauer Associates, Sunderland, MA, 2001).

24. G. Eisenman, H.R, Ionic selectivity revisited: the role of kinetic and equilibrium processes in ion permeation through channels. J. Membr. Biol. 76, 197–225 (1983).

25. O. A. Sineshchekov, E. G. Govorunova, H. Li, J. L. Spudich, Gating mechanisms of a natural anion channelrhodopsin. Proc. Natl. Acad. Sci. USA 112, 14236–14241 (2015).

26. J. Kuhne et al., Unifying photocycle model for light adaptation and temporal evolution of cation conductance in channelrhodopsin-2. Proc. Natl. Acad. Sci. USA 116, 9380–9389 (2019).

27. M. A. Dreier et al., Time-resolved spectroscopic and electrophysiological data reveal insights in the gating mechanism of anion channelrhodopsin. Commun Biol 4, 578 (2021).

28. M. K. Verhoefen et al., The photocycle of channelrhodopsin-2: Ultrafast reaction dynamics and subsequent reaction steps. Chemphyschem 11, 3113–3122 (2010).

29. O. A. Sineshchekov, E. G. Govorunova, J. Wang, H. Li, J. L. Spudich, Intramolecular proton transfer in channelrhodopsins. Biophys. J. 104, 807–817 (2013).

30. M. Baek et al., Accurate prediction of protein structures and interactions using a three-track neural network. Science, (2021).

31. B. Q. Minh et al., IQ-TREE 2: New models and efficient methods for phylogenetic inference in the genomic era. Mol. Biol. Evol. 37, 1530–1534 (2020).

32. D. T. Hoang, O. Chernomor, A. von Haeseler, B. Q. Minh, L. S. Vinh, UFBoot2: Improving the ultrafast bootstrap approximation. Mol. Biol. Evol. 35, 518–522 (2018).

33. I. Letunic, P. Bork, Interactive Tree Of Life (iTOL) v5: an online tool for phylogenetic tree display and annotation. Nucleic Acids Res. 49, W293–W296 (2021).

34. S. A. Waschuk, A. G. J. Bezerra, L. Shi, L. S. Brown, Leptosphaeria rhodopsin: Bacteriorhodopsin-like proton pump from a eukaryote. Proc. Natl. Acad. Sci. USA 102, 6879–6883 (2005).

35. M. Xue, B. V. Atallah, M. Scanziani, Equalizing excitation-inhibition ratios across visual cortical neurons. Nature 511, 596–600 (2014).

36. E. G. Govorunova et al., Cation and anion channelrhodopsins: Sequence motifs and taxonomic distribution. MBio 12, e0165621 (2021).

