## Supplemental Material for "Kalium rhodopsins: Natural light-gated potassium channels"

**This PDF file includes:**

Materials and Methods
Figs. S1 to S9
Table S1

### Materials and Methods

Bioinformatics and molecular biology

Initially, the predicted protein sequences encoded by the genes Hypho2016_00006030 and Hypho2016_00006031 (395 and 383 amino acid residues, respectively) were obtained from the database provided by reference (*21*) (https://www.ebi.ac.uk/biostudies/studies/S-BSST46), the file hyphochytrium_catenoides_predicted_proteins_renamed_modified. However, a large part of TM6 was missing from the Hypho2016_00006030 prediction. Therefore, we corrected its TM domain by performing TBLASTN search of the WGS data for *H. catenoides* strain ATCC 18719 (accession numbers FLMG00000000.1 and CAFC00000000.2) at the National Center for Biotechnology Information (NCBI), using the sequence provided (*21*) as a query. The resultant alignment allowed us to recover the full sequence (401 residues). DNA polynucleotides encoding the corrected transmembrane domains (residues 1-265) of Hypho2016_00006030 (*Hc*KCR2) and Hypho2016_00006031 (*Hc*KCR1) were optimized for human codon usage and synthesized at Genscript Biotech Corporation, (Piscataway, NJ). The sequence information was deposited to GenBank (accession numbers MZ826861 and MZ826862, respectively).

Rhodopsin sequences were aligned using MUSCLE as implemented in MegAlign Pro software v. 17.1.1 (DNASTAR Lasergene, Madison, WI) with default parameters. The transmembrane regions were predicted using RoseTTAFold modeling (*30*). Phylogeny was analyzed with IQ-TREE v. 2.1.2 (*31*) using automatic model selection and ultrafast bootstrap approximation (1000 replicates) (*32*). The best tree was visualized and annotated with iTOL v. 6.3 (*33*).

For expression in HEK293 (human embryonic kidney) cells the polynucleotides encoding the transmembrane domains of *Hc*KCR1 and *Hc*KCR2 were cloned into the mammalian expression vector pcDNA3.1 (Life Technologies, Grand Island, NY) in frame with an mCherry tag. For expression in *P. pastoris* the polynucleotide encoding the transmembrane domain of *Hc*KCR1 was fused with a C-terminal His7 tag and cloned into the pPICZαA vector (Invitrogen). For expression in mouse cortical neurons *Hc*KCR1 was tagged with EYFP (enhanced yellow fluorescent protein) at the C terminus and cloned into a pAAV-CAG vector.

HEK293 patch clamp electrophysiology

The cells were transfected using the ScreenFectA transfection reagent (Waco Chemicals USA, Richmond, VA). All-*trans*-retinal (Sigma) was added as a stock solution in ethanol at the final concentration of 5 µM. Measurements were performed 48-72 h after transfection with an Axopatch 200B amplifier (Molecular Devices, Union City, CA). The signals were digitized with a Digidata 1440A using pClamp 10 software (both from Molecular Devices). Patch pipettes with resistances of 2-5 MΩ were fabricated from borosilicate glass. The composition of solutions is shown in table S1. A 4 M salt bridge was used in all experiments. All *IE* dependencies were corrected for liquid junction potentials calculated using the ClampEx built-in LJP calculator (table S1). Continuous light pulses were provided by a Polychrome IV light source (T.I.L.L. Photonics GMBH, Grafelfing, Germany) in combination with a mechanical shutter (Uniblitz Model LS6, Vincent Associates, Rochester, NY; half-opening time 0.5 ms). Maximal quantum density at the focal plane of the 40× objective lens was ~7 mW $mm^{-2}$ at 540 nm. The action spectra were constructed by calculation of the initial slope of photocurrent in the linear range of the dependence on the quantum density (<25 µW $mm^{-2}$), corrected for the quantum density measured at each wavelength and normalized to the maximal value. Laser excitation was provided by a Minilite II Nd:YAG laser (532 nm, pulsewidth 6 ns, energy 12 mJ; Continuum, San Jose, CA). The current traces were logarithmically filtered using custom software. Curve fitting was performed by Origin Pro software (OriginLab Corporation, Northampton, MA). All measurements were carried out at room temperature (25° C).

Purification of *Hc*KCR1 from *Pichia pastoris*

The *Hc*KCR1-7His-pPICZαA plasmid was linearized with SacI and delivered into *P. pastoris* SMD1168 by electroporation. A single colony resistant to 0.5 mg/ml zeocin was picked and inoculated into buffered complex glycerol medium, after which the cells were transferred to buffered complex methanol (0.5%) medium supplemented with 5 µM all-*trans*-retinal (Sigma-Aldrich) and grown at 30°C with shaking at 200 rpm. After 24 h, the cells were harvested and disrupted in 100 ml of ice-cold buffer A (20 mM Hepes, pH 7.4, 150 mM NaCl, 1 mM EDTA, 5% glycerol) using a bead beater. Cell debris was removed by centrifugation at 5,000 × g for 10 min. Membrane fragments were collected by ultracentrifugation at 190,000 × g for 1 h, and then solubilized in 20 ml of buffer B (20 mM Hepes, pH 7.5, 350 mM NaCl, 5% glycerol) and 1% dodecyl-β-D-maltopyranoside (DDM) at 4°C for 1 h. Non-solubilized material was removed by ultracentrifugation at 110,000 × g for 1 h. The supernatant was mixed with nickel-nitrilotriacetic acid resin (Qiagen) with 15 mM imidazole and incubated at 4°C for 1 h. After washing the resin with buffer B containing 0.02% DDM and 40 mM imidazole, the protein was eluted with buffer B containing 0.02% DDM and 300 mM imidazole, concentrated using Amicon® Ultra centrifugal filters (Millipore) at 4°C, and washed with buffer B containing 0.02% DDM to remove imidazole.

UV-visual absorption spectroscopy and flash photolysis

Absorption spectra of purified *Hc*KCR1 were recorded using a Cary 4000 spectrophotometer (Varian, Palo Alto, CA). Light-induced absorption changes were measured with a laboratory-constructed crossbeam apparatus. Excitation flashes (532 nm, 6 ns, 12 mJ) were provided by a Minilite II Nd:YAG laser (Continuum, San Jose, CA). Measuring light was from a 250-W incandescent tungsten lamp combined with a McPherson monochromator (model 272, Acton, MA). Absorption changes were detected with a Hamamatsu Photonics (Bridgewater, NJ) photomultiplier tube (model R928), protected from excitation laser flashes by a second monochromator of the same type. Signals were amplified by a low noise current amplifier (model SR445A; Stanford Research Systems, Sunnyvale, CA) and digitized with a GaGe Octopus digitizer board (model CS8327, DynamicSignals LLC, Lockport, IL), maximum sampling rate 50 MHz. Logarithmic filtration of the data was performed using the GageCon program (*34*).

Mice

All procedures to maintain and use mice were approved by the Institutional Animal Care and Use Committee at Baylor College of Medicine. Mice were maintained on a 14 hr:10 hr light:dark cycle with regular mouse chow and water *ad libitum*. Experiments were performed during the light cycle. ICR (CD-1) female mice were purchased from Baylor College of Medicine Center for Comparative Medicine. C57BL6/J male mice were obtained from Jackson Laboratory (stock numbers 000664). Both male and female mice were used in the experiments.

*In utero* electroporation

Female ICR mice were crossed with male C57BL6/J to obtain timed pregnancies. *In utero* electroporation was performed as previously described (*35*) with a square-wave pulse generator (Gemini X2, BTX Harvard Bioscience). To express *Hc*KCR1 in the layer 2/3 pyramidal neurons of the somatosensory cortex, pAAV-CAG-*Hc*KCR1-EYFP (2.5 µg/µl as final concentration) and pCAG-tdTomato (0.1 µg/µl as final concentration) was injected into the lateral ventricle at embryonic day 14.5 or 15, followed by electroporation. Transfected pups were identified by the transcranial fluorescence of tdTomato with a MZ10F stereomicroscope (Leica) 1 day after birth.

Brain slice electrophysiology

Mice were used at the age of 2–4 weeks for acute brain slice electrophysiology experiments. Mice were anesthetized by an intraperitoneal injection of a ketamine and xylazine mix (80 mg/kg and 16 mg/kg, respectively) and transcardially perfused with cold (0–4°C) slice cutting solution containing 80 mM NaCl, 2.5 mM KCl, 1.3 mM $NaH_2PO_4$, 26 mM $NaHCO_3$, 4 mM $MgCl_2$, 0.5 mM $CaCl_2$, 20 mM D-glucose, 75 mM sucrose and 0.5 mM sodium ascorbate (315 mOsmol, pH 7.4, saturated with 95% O2/5% CO2). Brains were removed and sectioned in the cutting solution with a VT1200S vibratome (Leica) to obtain 300 μm coronal slices. Slices were incubated in a custom-made interface holding chamber containing slice cutting solution saturated with 95% O2/5% CO2 at 34°C for 30 min and then at room temperature for 20 min to 10 hr until they were transferred to the recording chamber.

Recordings were performed on submerged slices in artificial cerebrospinal fluid (ACSF) containing 119 mM NaCl, 2.5 mM KCl, 1.3 mM $NaH_2PO_4$, 26 mM $NaHCO_3$, 1.3 mM $MgCl_2$, 2.5 mM $CaCl_2$, 20 mM D-glucose and 0.5 mM sodium ascorbate (305 mOsmol, pH 7.4, saturated with 95% O2/5% CO2, perfused at 3 ml/min) at 30–32°C. For whole-cell recordings, a $K^+$-based pipette solution containing 142 mM $K^+$-gluconate, 10 mM HEPES, 1 mM EGTA, 2.5 mM $MgCl_2$, 4 mM ATP-Mg, 0.3 mM GTP-Na, 10 mM $Na_2$-phosphocreatine (295 mOsmol, pH 7.35) was used. Membrane potentials reported in Fig. 3 were not corrected for liquid junction potential that was experimentally measured as 12.5 mV.

Neurons were visualized with video-assisted infrared differential interference contrast imaging, and fluorescent neurons were identified by epifluorescence imaging under a water immersion objective (40×, 0.8 numerical aperture) on an upright SliceScope Pro 1000 microscope (Scientifica) with an infrared IR-1000 CCD camera (DAGE-MTI). Data were low-pass filtered at 4 kHz and acquired at 10 kHz with an Axon Multiclamp 700B amplifier and an Axon Digidata 1440A Data Acquisition System under the control of Clampex 10.7 (Molecular Devices). Data were analyzed offline using Clampfit (Molecular Devices). For photostimulation of *Hc*KCR1-expressing neurons, green light was emitted from a collimated light-emitting diode (LED) of 565 nm (Thorlabs M565L3). The LEDs were driven by a LED driver (Throlabs LEDD1B) under the control of an Axon Digidata 1440A Data Acquisition System and Clampex 10.7. Light was delivered through the reflected light fluorescence illuminator port and the 40× objective.

Photocurrents were recorded by whole-cell voltage clamp in response to 1-s 565 nm light stimulation (4.5 mW $mm^{-2}$). Only recordings with series resistance below 20 MΩ were included. To test the recovery of photocurrents in the dark, *Hc*KCR1-expressing neurons were held at -45 mV and stimulated with various inter-stimulus intervals (ISI). To test current-voltage relationship, photocurrents were recorded at membrane voltages from -85 to -45 mV with 5-mV steps and 30-s ISI. Action potentials of *Hc*KCR1-expressing neurons were evoked by injecting a series of 1.5-s depolarizing current pulses (0.1–0.5 nA) in whole-cell current clamp mode. 1-s 565 nm light stimulation (4.5 $mW/mm^2$) was applied in the middle of current injections with 30-s ISI. Light stimulation and control trials were interleaved.

Fluorescent imaging

After electrophysiology recordings, brain slices were fixed overnight in 4% paraformaldehyde in PBS (pH 7.4), cryoprotected with 30% sucrose in PBS, and frozen in optimum cutting-temperature medium until sectioning. A HM 450 Sliding Microtome (Thermo

Scientific) was used to further section the slices to obtain 50 µm slices. Images were acquired on an Axio Zoom.V16 Fluorescence Stereo Zoom Microscope (Zeiss) and processed using Matlab (MathWorks).

Statistics

For experiments in HEK293 cells, descriptive statistics was calculated by Origin software. The data are presented as mean ± sd or sem values, as indicated in the figure legends; the data from individual replicates are also shown when appropriate. The sample size was estimated from previous experience and published work on similar subjects, as recommended by the NIH guidelines. No normal distribution of the data was assumed; when a specific statistics hypothesis was tested, the non-parametric Mann-Whitney test (implemented in Origin software) was used.

For experiments in neurons, statistical analyses were performed with Prism 9 (GraphPad Software). Paired *t* test was used in Fig. 3D and Fig. 3G, Wilcoxon matched-pairs signed rank test in Fig. 3C, and Multiple Wilcoxon matched-pairs signed rank test in Fig. 3I. All statistical tests were two-tailed with an alpha of 0.05. All reported sample numbers (n) represent biological replicates that are the numbers of recorded neurons.

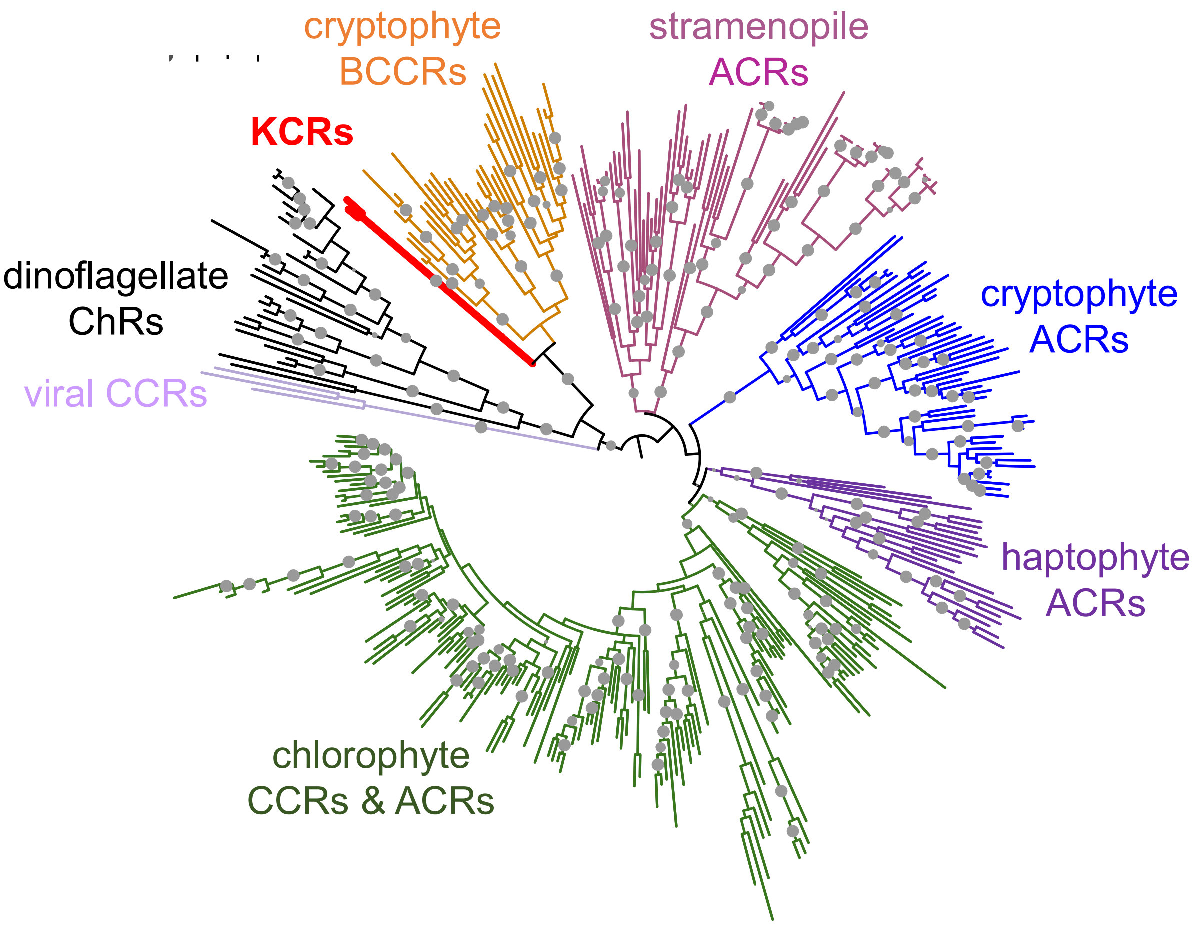


**Fig. S1. Phylogenetic relations of KCRs with other known ChRs.**

The branches are colored to distinguish different ChR families. The leaves corresponding to the KCRs characterized in this study are shown as thick red lines. A full list of other ChR sequences used to create the tree can be found in reference (*36*). The gray circles show ultrafast bootstrap support values above 95%.

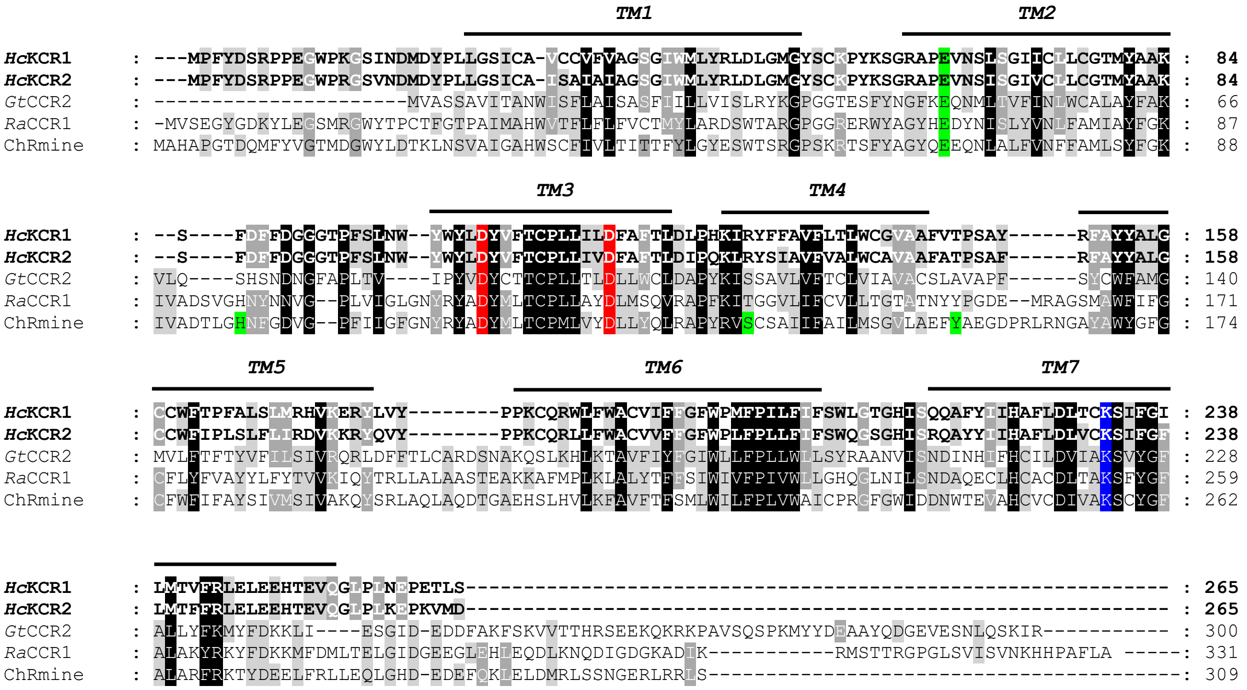


**Fig. S2. The alignment of KCRs and representative cryptophyte BCCRs.**

The black lines show the predicted transmembrane helices (TM1-TM7). The Schiff base lysine is highlighted blue, the conserved aspartates corresponding to Asp85 and Asp96 of bacteriorhodopsin, red, and the residues implicated in trimer formation in ChRmine, green. *Gt*CCR2, *Guillardia theta* CCR2; *Ra*CCR1, *Rhodomonas abbreviata* CCR1.

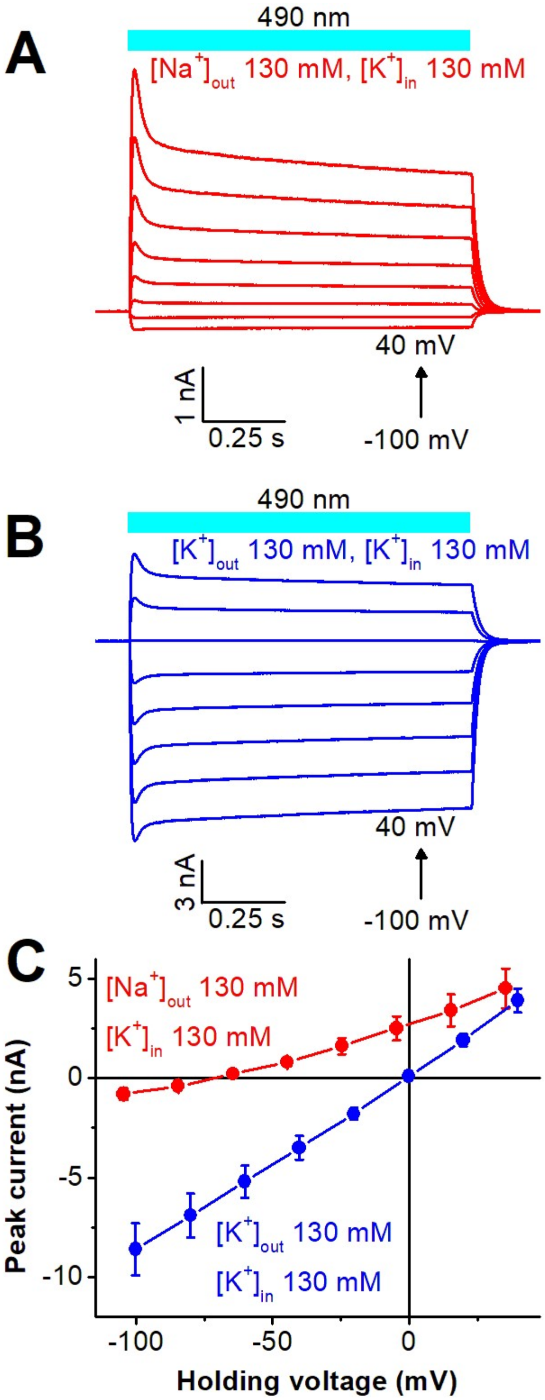


**Fig. S3. Electrophysiological characterization of *Hc*KCR2.**

(**A** and **B**) Series of *Hc*KCR2 photocurrents recorded in response to 1-s light pulses under indicated ionic and voltage conditions. (**C**) The *IE* curves measured under indicated ionic conditions (mean ± sem, n = 7 cells).

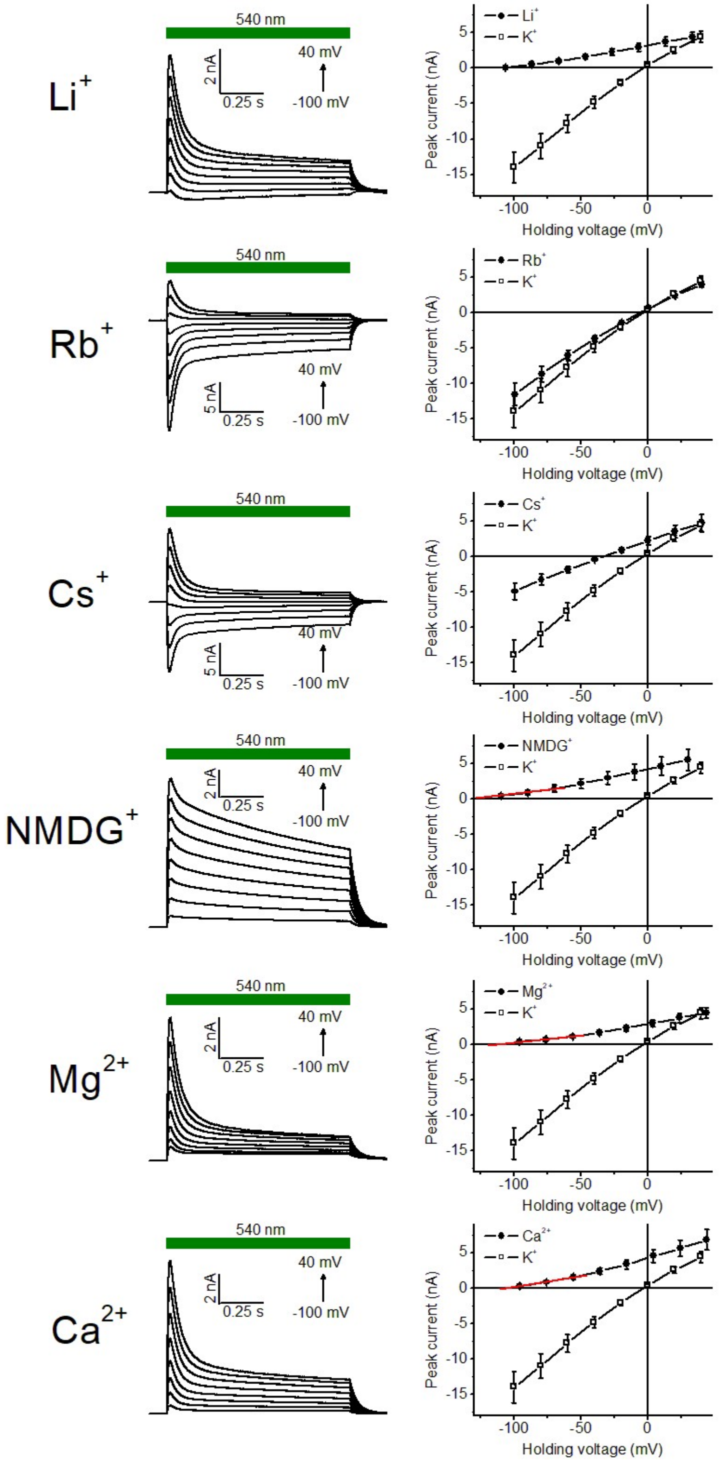


**Fig. S4. Analysis of relative permeabilities of *Hc*KCR1 for metal cations and $NMDG^+$.**

Left, *Hc*KCR1 photocurrents recorded in response to 1-s light pulses with 130 mM KCl in the pipette and 130 mM of the indicated ion in the bath. Right, the corresponding *IE* curves (mean ± sem, n = 7 cells). The red lines show linear approximations used to determine the $E_{rev}$.

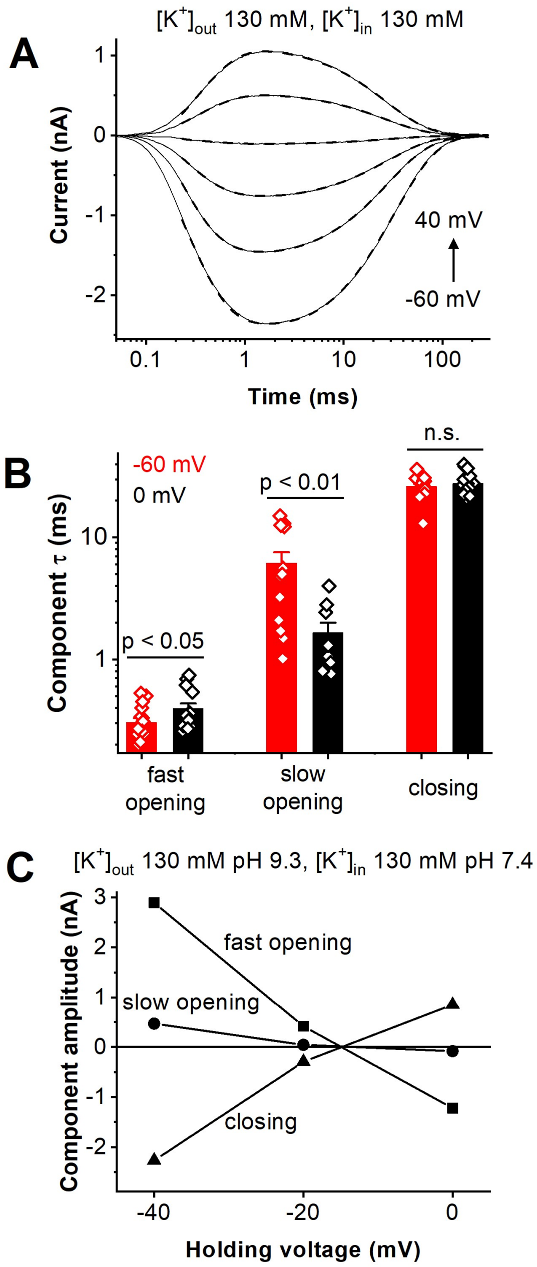


**Fig. S5. pH and voltage dependence of photocurrent kinetic components.**

(**A**) Photocurrent traces (thin solid lines) recorded from *Hc*KCR1 in response to 6-ns laser flashes at 20-mV voltage increments under indicated ionic conditions and their multiexponential approximations (dashed lines). (**B**) The time constants (τ) of the three kinetic components of channel currents at -60 (red) and 0 (black) mV (mean ± sem). The data for individual cells are shown as diamonds. The P values were calculated by the Mann-Whitney test. n.s., not significant. (**C**) The voltage dependence of the three kinetic components of channel currents.

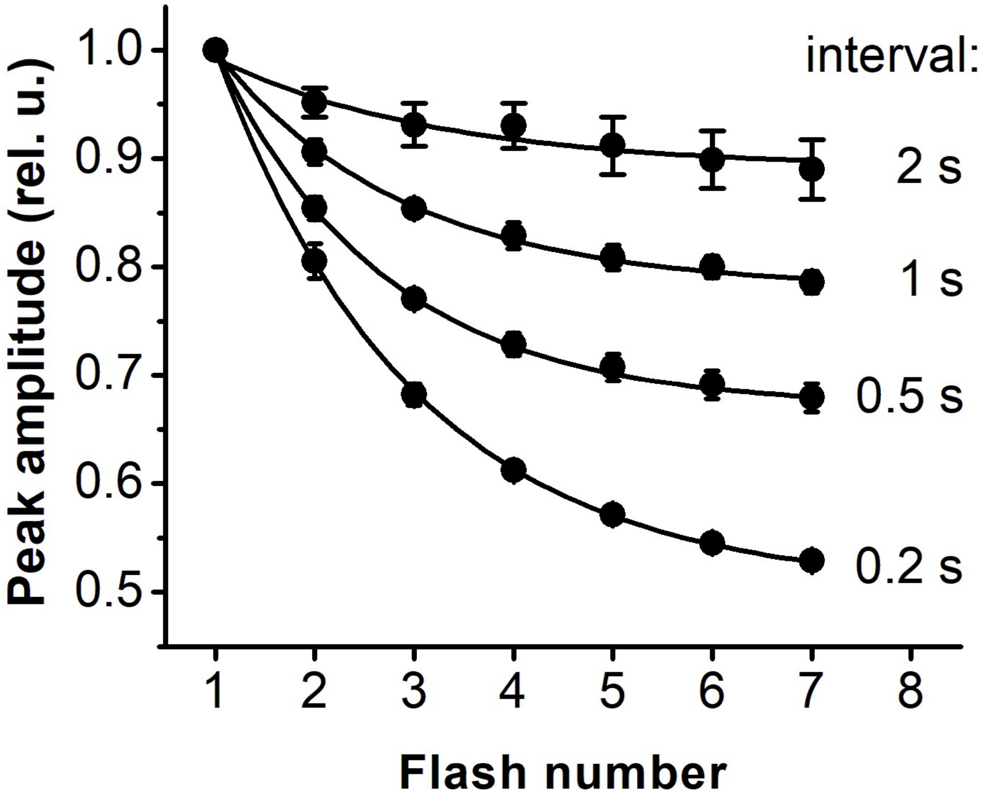


**Fig. S6. Peak amplitude channel currents recorded at varied time intervals between laser flashes.**

The data points are mean ± sem, n = 5 cells.

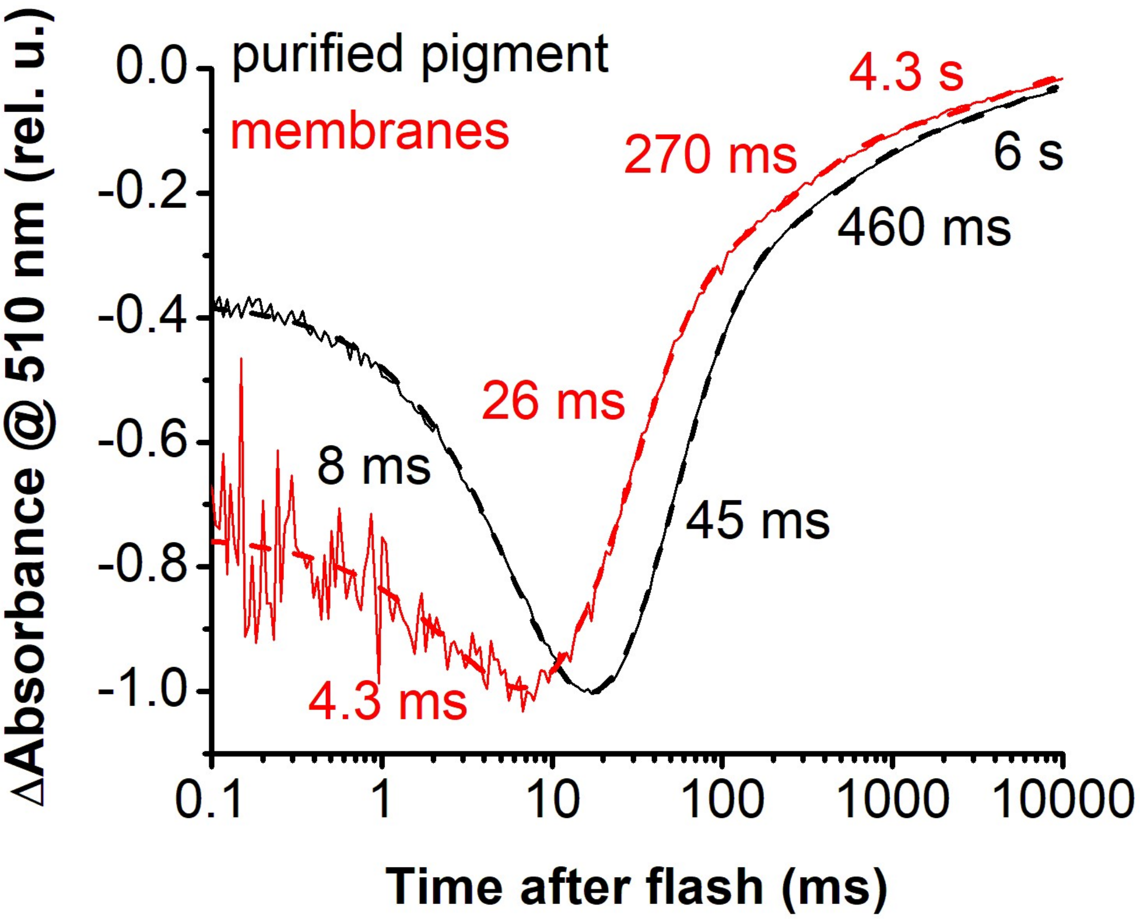


**Fig. S7. Laser flash-induced absorption changes of *Hc*KCR1 in detergent (black) and *Pichia* membranes (red).**

Experimental data are shown as thin solid lines, and their multiexponential approximations, as dotted lines.

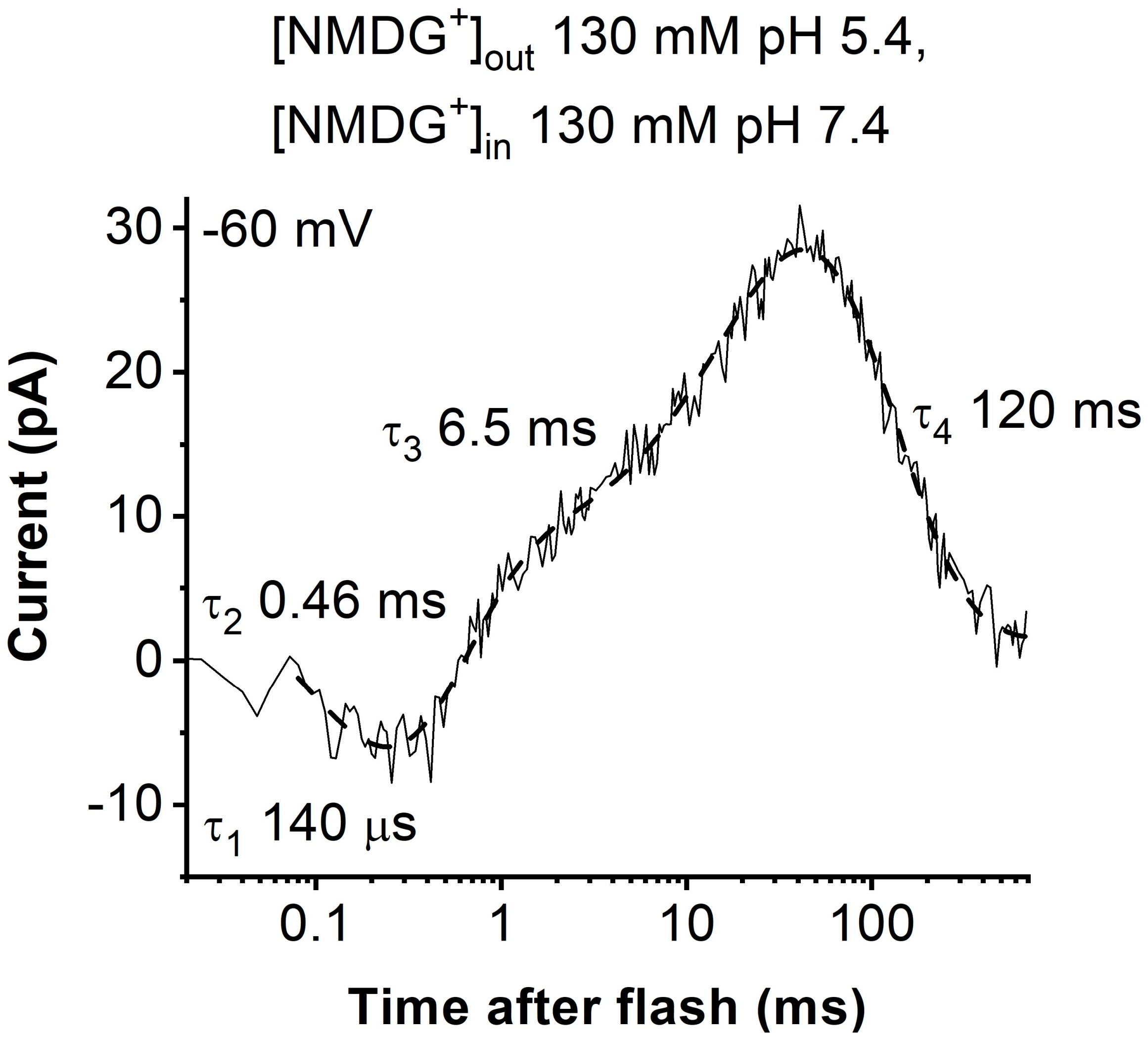


**Fig. S8.** ***Hc*KCR1 photocurrent traces in the absence of permeant metal cations at bath pH 5.4.**

Experimental data are shown as thin solid lines, and their multiexponential approximations, as dotted lines.

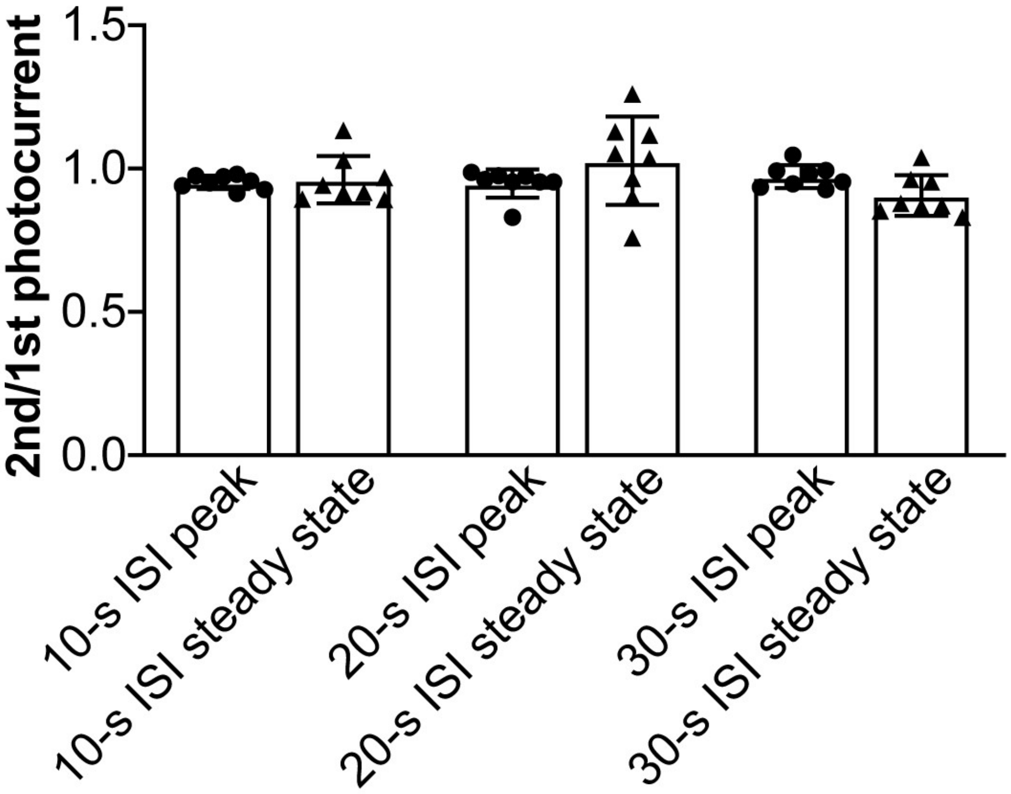


**Fig. S9. *Hc*KCR1 photocurrent recovery in the dark in experiments with pulses of continuous light.**

The ratio of the peak or steady-state currents evoked by two 1-s light pulses applied with a 10-s, 20-s, or 30-s interval. ISI, inter-stimulus interval. Data are expressed as mean ± sem; n = 8 neurons.

**Table S1.**

| | NaCl | KCl | LiCl | RbCl | CsCl | NMDG | $MgCl_2$ | $CaCl_2$ | HEPES | Glucose | LJP pip. stand |
|---|---|---|---|---|---|---|---|---|---|---|---|
| **Pipette standard** | — | 130 | — | — | — | — | 2 | — | 10 | — | — |
| **Pipette $NMDG^+$** | — | — | — | — | — | 130 | 2 | — | 10 | — | — |
| **Bath $Na^+$** | 130 | — | — | — | — | — | 2 | 2 | 10 | 10 | 4.4 |
| **Bath $K^+$** | — | 130 | — | — | — | — | 2 | 2 | 10 | 10 | 0.2 |
| **Bath $Li^+$** | — | — | 130 | — | — | — | 2 | 2 | 10 | 10 | 6.7 |
| **Bath $Rb^+$** | — | — | — | 130 | — | — | 2 | 2 | 10 | 10 | -0.5 |
| **Bath $Cs^+$** | — | — | — | — | 130 | — | 2 | 2 | 10 | 10 | -0.4 |
| **Bath $NMDG^+$** | — | — | — | — | — | 130 | 2 | 2 | 10 | 10 | 9.9 |
| **Bath $Mg^{2+}$** | — | — | — | — | — | — | 65 | 2 | 10 | 10 | -3.8 |
| **Bath $Ca^{2+}$** | — | — | — | — | — | — | 2 | 65 | 10 | 10 | -4.2 |

Composition of pipette and bath solutions and liquid junction potentials in experiments with HEK293 cells. Abbreviations: HEPES, 4-(2-hydroxyethyl)-1-piperazineethanesulfonic acid; LJP, liquid junction potential; NMDG, N-Methyl-D-glucamine. All concentrations are in mM.
